# Estimating the quality of eukaryotic genomes recovered from metagenomic analysis

**DOI:** 10.1101/2019.12.19.882753

**Authors:** Paul Saary, Alex L. Mitchell, Robert D. Finn

## Abstract

Eukaryotes make up a large fraction of microbial biodiversity. However, the field of metagenomics has been heavily biased towards the study of just the prokaryotic fraction. This focus has driven the necessary methodological developments to enable the recovery of prokaryotic genomes from metagenomes, which has reliably yielded genomes from thousands of novel species. More recently, microbial eukaryotes have gained more attention, but there is yet to be a parallel explosion in the number of eukaryotic genomes recovered from metagenomic samples. One of the current deficiencies is the lack of a universally applicable and reliable tool for the estimation of eukaryote genome quality. To address this need, we have developed EukCC, a tool for estimating the quality of eukaryotic genomes based on the dynamic selection of single copy marker gene sets, with the aim of applying it to metagenomics datasets. We demonstrate that our method outperforms current genome quality estimators and have applied EukCC to datasets from two different biomes to enable the identification of novel genomes, including a eukaryote found on the human skin and a *Bathycoccus* species obtained from a marine sample.

## Introduction

The past two decades have seen a dramatic advancement in our understanding of the microscopic organisms present in environments (known as microbiomes) such as oceans, soil and host-associated sites, like the human gut. Most of this knowledge has come from the application of modern DNA sequencing techniques to the collective genetic material of the microorganisms, using methods such as metabarcoding (amplification of marker genes) or metagenomics (shotgun sequencing). Based on the analysis of such sequence data, it is thought that up to 99% of all microorganisms are yet to be cultured (Rinke et al. 2013).

To date, the overwhelming number of metabarcoding and metagenomics studies have focused on the bacteria that are present within a sample. However, viruses and eukaryotes are also important members of the microbial community, both in terms of number and function (Paez-Espino et al. 2016; Carradec et al. 2018; Olm et al. 2019; Karin et al. 2019). Indeed, the unicellular protists and fungi are estimated to account for about ~17% of the global microbial biomass. Within the microbial eukaryotic biomass, the genetically diverse unicellular organisms known as protists account for as much as ~25% (Bar-On et al. 2018). Today, the increasing number of completed genomes has revealed that the “protists” classification encompasses a number of divergent sub-clades: animals and some protists are encompassed in the Opisthokonta clade; others are grouped into the Amoebozoa or (T)SAR ((Telonemids), Stramenopiles, Alveolates, and Rhizaria) clades, or into further groups. However, the exact root of the overall eukaryotic tree and the number of primary clades remains a topic of discussion (Baldauf 2003; Burki 2014; Burki et al. 2019).

Despite the increase in the number of complete and near complete genomes, metabarcoding and metagenomic approaches that have included the analysis of microbial eukaryotes have demonstrated that the true diversity of protists is far greater than that currently reflected in the genomic reference databases (such as Ref-Seq or ENA). For example, a recent estimate based on metabarcoding sequencing suggests that 150,000 eukaryotic species exist in the oceans alone (Vargas et al. 2015), but only 4,551 representative species have an entry in GenBank (15. Nov 2019). Thus, if the functional role of a microbiome is to be completely understood, we need to know what these as yet uncharacterised organisms are and the functional roles they are performing.

Currently, one of the best approaches for understanding microbiome function is through the assembly of shotgun reads (usually 200-500 bp long) to obtain longer contigs (typically in the range of 2000-500,000 bp). These contigs provide access to complete proteins, which may then be interpreted within the context of surrounding genes. In the last few years, it has become commonplace to extend this type of analysis to recover putative genomes, termed metagenome assembled genomes (MAGs). MAGs are generated by grouping contigs into sets that are believed to have come from a single organism – a process known as binning. However, even after binning MAGs, they vary in their completeness and can be fragmented, due to a combination of biological (e.g. abundance of microbes), experimental (e.g. depth of sequencing) and technical (e.g. algorithmic) reasons. Furthermore, the computational methods used for binning the contigs can sometimes fail to distinguish between contigs that have come from different organisms, leading to a chimeric genome (termed contamination). As highlighted above, reference databases are incomplete, so estimating the quality of a MAG in terms of completeness and contamination can not rely on genomic comparisons. In the absence of a reference genome, quality estimates for MAGs have used universal single copy marker genes (SCMGs) (Parra et al. 2007; Mende et al. 2013; Simão et al. 2015; Parks et al. 2015). As these genes are expected to only occur once within a genome, comparing the number of SCMGs found within a binned genome to the number of expected marker genes provides an estimation of completeness, while additional copies of a marker gene can be used as an indicator of contamination. After such evaluations, binned genomes achieving a certain quality can be classified as either medium or high quality (Bowers et al. 2017).

Due to biases in sampling and extraction methods, the majority of MAGs produced to date correspond to prokaryotic organisms. For prokaryotic MAGs, CheckM (Parks et al. 2015) is the most widely used tool to estimate completeness and contamination, although other approaches have also been used (Pasolli et al. 2019) and individual sets of prokaryotic SCMGs are also provided by BUSCO (Simão et al. 2015; Waterhouse et al. 2018) as well as by anvi’o (Eren et al. 2015). However, even with size fractionation of samples to enrich for prokaryotes prior to library preparation, eukaryotic cells frequently remain in the samples, with some eukaryotic DNA recovered as MAGs (Delmont et al. 2018; West et al. 2018), while others use size fractionation specifically to enrich for eukaryotes(Karsenti et al. 2011; Carradec et al. 2018).

As with the quality estimation of bacterial MAGs, SCMGs have been used to assess eukaryotic isolate genomes. CEGMA (Parra et al. 2007) used 240 universal single copy marker genes identified from six model organisms to estimate genome completeness, which was then superseded by BUSCO. The major advance of BUSCO compared to CEGMA, was the provision of curated sets of marker genes for several eukaryotic and prokaryotic clades, in addition to the single universal eukaryotic marker gene set. While BUSCO (version 3) provides sets to estimate completeness of eukaryota, protists, plants and fungi, it remains up to the user to select which is the most suitable set when assessing genome quality. Although BUSCO has been used for quality metric calculation of eukaryotic MAGs (West et al. 2018), this manual selection can be challenging, especially when dealing with large numbers of genomes from unknown species. Besides these more universal approaches, others have focused on certain clades of Eukaryotes: FGMP (Cissé and Stajich 2019) estimates genome quality of Fungal isolate genomes for which it utilises both SCMGs and also highly conserved regions found within fungal genomes. For protists, anvi’o provides a reduced number of profiles from the BUSCO ‘protist’ set to estimate the genome quality of eukaryotic MAGs (unpublished).

Here we investigate the performance of current approaches across different eukaryotic clades and describe EukCC, an unsupervised method for the estimation of eukaryotic genome quality in terms of completion and contamination, with a particular view of applying this tool to eukaryotic MAGs.

## Results

### Evaluation of BUSCO across different eukaryotic clades

To determine the applicability of BUSCO for evaluating the quality of eukaryotic MAGs, we first tested how the more general eukaryotic BUSCO set performed in terms of assessing the completeness and contamination for a range of eukaryotic isolate genomes. Briefly, fungal and protist genomes were downloaded from the NCBI Reference Sequence Database (RefSeq) and estimated using BUSCO in ‘*genome mode*’, which employs AUGUSTUS for gene prediction (Keller et al. 2011), with the eukaryota SCMG set (‘eukaryota_odb9’). Fungi and protist genomes were additionally estimated using the fungal (‘fungi_odb9’) and protist (‘protists_ensembl’) set respectively. As these genomes are of high quality and manually curated, it was anticipated that they should have very high levels of completeness and minimal levels of contamination.

To understand the overlap between the eukaryotic BUSCO set and the selected genomes, we counted the number of matched BUSCOs in each taxonomic clade containing at least 3 reference genomes. While BUSCO reports complete, fragmented and duplicated BUSCOs, for the sake of simplicity we summarized all these as ‘matched’ BUSCOs (Figure 1 A). One of the main applications of BUSCO has been the assessment of fungal genomes, which also represent the most numerous eukaryotic genomes in the reference databases. Thus, it was unsurprising that > 95% of the 303 eukaryotic BUSCOs were matched in genomes coming from Ascomycota, Mucoromycota and Basidiomycota. However, BUSCO performed less well on eukaryotic genomes arising from other taxonomic groups. Notably, the numbers of BUSCOs found in Amoebozoa genomes varied greatly, with a median of 88.78%, but ranging between 69.6% for Entamoebidae (number of species, n = 4) to 94.9% for the four further Amoebozoa families (n = 6). More surprising was that the Ciliophora genomes (n = 4) rarely matched BUSCO eukaryotic marker genes, with a median of 1.16% of BUSCOs matched.

**Figure 1:**
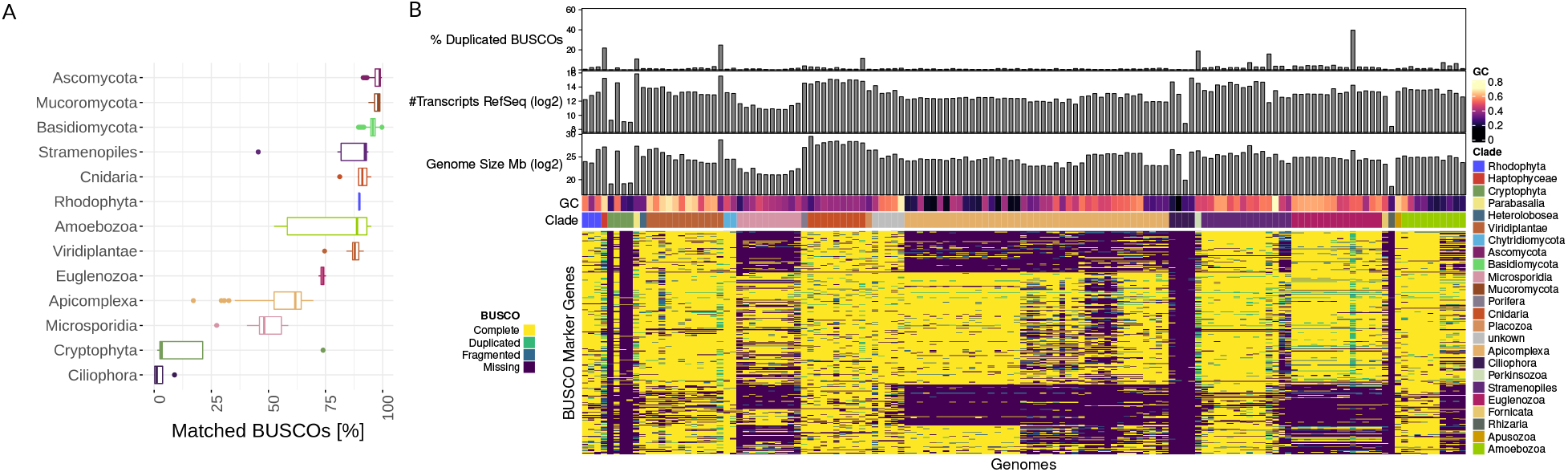
**A)** We downloaded eukaryotic RefSeq genomes excluding bilateria and vascular plants, and ran BUSCO in ‘*genome mode*’ using the ‘eukaryota_odb9’ set. For each clade we summarized the number of BUSCO markers matched. For Fungal clades, such as Ascomycota, Mucoromycota and Basidiomycota, most BUSCOs matched a single target – suggesting 100% completeness of the reference genomes. However, in other clades a substantial fraction of BUSCOs were frequently not matched (Apicomplexa, for example). **B)** For species not belonging to Fungal clades, we created a matrix using the detailed BUSCO results. Genomes are sorted taxonomically (using the assigned NCBI taxonomy) in columns and the result for each BUSCO in rows. The matrix is coloured according to the BUSCO result, which reports complete, duplicated, fragmented and missing marker genes. Fragmented hits are reported if only part of the BUSCO was detected. Above is shown the percentage of duplicated BUSCOs, the number of the RefSeq transcripts for each genome, the genome size and the GC content. In some clades, there is a clear relationship between the genome taxonomy and missing and BUSCOs. In the case of Micosporida and Apicomplexa, but also for Euglenozoa, this relationship is especially strong.

We also evaluated the BUSCO protist set in the same way. Somewhat counterintuitively, using this more specific set the mean proportion of matched BUSCOs in Amoebozoa dropped from 88.78% to 78.37%, yet increased for Apicom-plexa from 61.72% to 68.37%. In other taxa, such as Stramenopiles, the range of missing BUS-COs increased (Figure S1). This suggests that the use of a more specific BUSCO set can improve predictions, but does not resolve the problem of inaccurate estimation of completeness in specific clades.

To determine if the underestimation in clades other than fungi is random or caused by systematic biases, we created a matrix containing all found, missing, fragmented or complete BUS-COs in all analysed reference genomes, excluding Basidiomycota, Mucoromycota and Ascomy-cota (Figure 1 B, see Methods). We arranged the columns based on the NCBI taxonomy and rows using *k*-modes clustering. Within certain clades, such as Cryptophyta, Micosporida and Apicomplexa, the same BUSCOs were often missing across a large number of species. For each BUSCO, we evaluated whether it was missing in at least half the species of a given clade. Subsequently, when disregarding any BUSCO missing in at least three clades, the number of BUSCOs in the eukaryota set was reduced from 303 to 86.

Taken together, this shows that the BUSCO eukaryota set does not perform uniformly across all eukaryotic clades. Others have observed similar issues when investigating individual species or clades (Benites et al. 2019; Hackl et al. 2019). We also investigated whether factors, such as genome size, GC content or proteome size, could account for the bias in matching BUSCOs, but taxonomic lineage represented the single strongest signal.

### Influence of Gene Prediction on BUSCO matches

To understand whether issues with *de novo* gene prediction could be the cause of the missing BUSCO matches, we additionally ran BUSCO in ‘*protein mode*’ on the genome protein annotations provided by RefSeq and proteins predicted using GeneMark-ES (Ter-Hovhannisyan et al. 2008; Figure S1 C). When running BUSCO in this mode against RefSeq protein annotations, the number of matched BUSCOs increased overall, indicating that *de novo* prediction methods do account for some of the loss of sensitivity. However, the general pattern of missing markers across clades remained. Taking Ciliophora as an extreme example, the median of matched markers was 1.2% in ‘*genome mode*’, which was increased to 76.2% using RefSeq annotations. For other clades the differences were less substantial but still observable. For example, in Apicomplexa 61.7% of BUSCOs were matched using AUGUSTUS, rising to 73.9% using GeneMark-ES and 74.2% with RefSeq annotations. Notably, GeneMark-ES failed to run on several genomes of the Cryptophyta and Ciliophora clades, as well as for the single Rhizaria genome, which BUSCO estimated in ‘*genome mode*’ to have close to 100% missing markers. The primary reason GeneMark-ES did not work for a genome was a lack of suitable training data: out of six failed annotation attempts, five had four or less contigs included in the training phase of GeneMark-ES.

### Establishing specific single copy marker gene libraries

To more accurately compute quality estimates for novel genomes, we wanted to define sets of SCMGs that were comprehensive for microbial eukaryotes, as well as being both sensitive and specific. As shown above, BUSCO produces sets of SCMGs for specific clades which can be more precise in quality estimation. Building on this observation, we aimed at defining multiple sets of SCMGs covering a large range of protists and fungi. We anticipate that a key use case of the marker gene library will be the application of it to poorly characterised genomes, and as such, the genes are likely to be identified by *de novo* prediction. We therefore chose to (re-)annotate all eukaryotic RefSeq species (not belonging to bilateria or vascular plants) using GeneMark-ES generated gene predictions to closely represent the use case. In addition, this ensures that only proteins predicted using *de novo* approaches are used as marker genes. GeneMark-ES was applied, as we previously demonstrated that the tool works well across a large range of species and generally performs closer to the RefSeq annotation benchmark. Additionally, we added all species that are used as references in UniProtKB. The resulting proteins were then annotated with the family-level profile HMMs from PANTHER 14.1 using hmmer (version 3.2). We choose PANTHER, as among tested databases it has been shown to have the largest coverage of the analysed proteins (Mitchell et al. 2019), and because the PANTHER profile HMMs model full-length protein families rather than their constituent globular domains.

In order to increase paralog separation and minimise local matches caused by common domains, we aimed to define profile specific bit score thresholds. To achieve this, we relied on a taxonomically balanced set of species, across which, for each profile we identified the bit score threshold leading to the highest number of single copy matches (see Methods).

Thereafter, to define clade specific SCMGs, we first constructed a reference tree for the given genomes using 55 widely occurring SCMGs (from here on termed “reference set”) (see Methods). In each clade of the tree we checked for SCMGs with a prevalence of at least 98%. A set of marker genes was then defined whenever we found 20 or more PANTHER families in a clade matching the aforementioned prevalence threshold. Using this approach, we were able to define 477 SCMG sets across the entire tree. In contrast to BUSCO and CEGMA, we were not able to identify SCMGs applicable to the entire eukaryotic kingdom, but found sets applicable to many subclades. While this is desirable for specificity, the obvious drawback is knowing which is the most appropriate set to use – it would be impractical to manually assign the most appropriate set (especially if a large number of different genomes were to be assessed). Thus, we developed EukCC, a software package to select the most appropriate SCMGs, and use these to estimate genome quality.

### Automatically selecting the appropriate single copy marker gene set

To select the most specific set of SCMGs for a novel genome of unknown taxonomic lineage, EukCC performs an initial taxonomic classification by annotating the *de novo* predicted proteins using the 55 widely occurring SCMGs reference set. Pplacer (Matsen et al. 2010) is then applied to phylogenetically contextualise each match within the reference tree. Tracing each placement in the tree, EukCC determines the lowest common ancestor (LCA) node for which an SCMG set is defined in the database.

As may be expected, while pplacer often places all sequences in a simple, narrow region of the reference tree, occasional placements occur within inconsistent, distantly related clades. In such cases, no single set of SCMGs may encompass all locations. To overcome this, in these cases, the SCMG set that encapsulates the largest fraction of the placements is located. While this process overcomes cases where outlying placements occur due to incorrect or inconsistent placements, this approach may select an incorrect SCMG set if the matches to the reference SCMGs from a novel genome can not reliably be placed in the tree. To help control for this, EukCC always reports how many profiles are covered in a set and provides the option of plotting the placement locations (Figure S5). Thus, in a situation where a set was chosen that only encompasses a fraction of the reference SCMGs, a more in-depth analysis of this MAG could, and should, be carried out.

After the initial placement, EukCC assesses the completeness and contamination in a second step by annotating all proteins with the profiles that are expected to be single copy within the assigned clade. EukCC then reports the fraction of single copy markers found and the fraction of duplicated marker genes, corresponding to the completeness and contamination score, as provided for prokaryotes by CheckM. Additionally, EukCC uses the inferred placement to give a simple phylogenetic lineage estimation based on the consensus NCBI taxonomy of the species used to construct the chosen evaluation set.

### Comparison of BUSCO to EukCC quality estimates

Having established new sets of SGMGs and having developed EukCC for their selection, we next evaluated the accuracy of our approach for estimating completeness and contamination. To do so, we used both EukCC and BUSCO to estimate the completeness and contamination of 21 RefSeq genomes, from 7 different clades, that were not used to establish the EukCC SCMGs. As these were complete genomes, we simulated varying amounts of completeness and contamination (see Methods). Furthermore, to make the comparison balanced, we used the taxonomy assigned to each genome to select the most specific BUSCO sets, while letting the EukCC algorithm dynamically select the SCMGs set from our library of clade specific SCMGs. As we showed earlier that the *de novo* gene prediction can have an influence on the BUSCO results, we ran BUSCO using AUGUSTUS as well as in ‘*protein mode*’ on GeneMark-ES predicted proteins, which are also used by EukCC.

When estimating completeness across simulated genomes with no added contamination, EukCC performed better than BUSCO using either AUGUSTUS or GeneMark-ES. BUSCOs estimates for simulated genomes with more than 95% completeness and no contamination were better when relying on GeneMark-ES, but underestimated completeness with a median of 21.0% compared to 2.5% for EukCC (Figure 2 A). It is worth noting that while the completeness estimates of BUSCO can deviate strongly from the expected value, the degree of error varied across different taxonomic groups. For example within fungi, estimates deviate below 2% from the expected value (Figure S2). Across all tested clades EukCCs completeness estimates are closer to the expected value than BUSCOs. EukCC performed best for fungi and Alveolates, but underestimates completeness for simulated genomes (≥ 95% completeness, ≤ 5% contamination) of Amoebozoa and Viridiplantae by 14.2% and 7.7% respectively.

**Figure 2:**
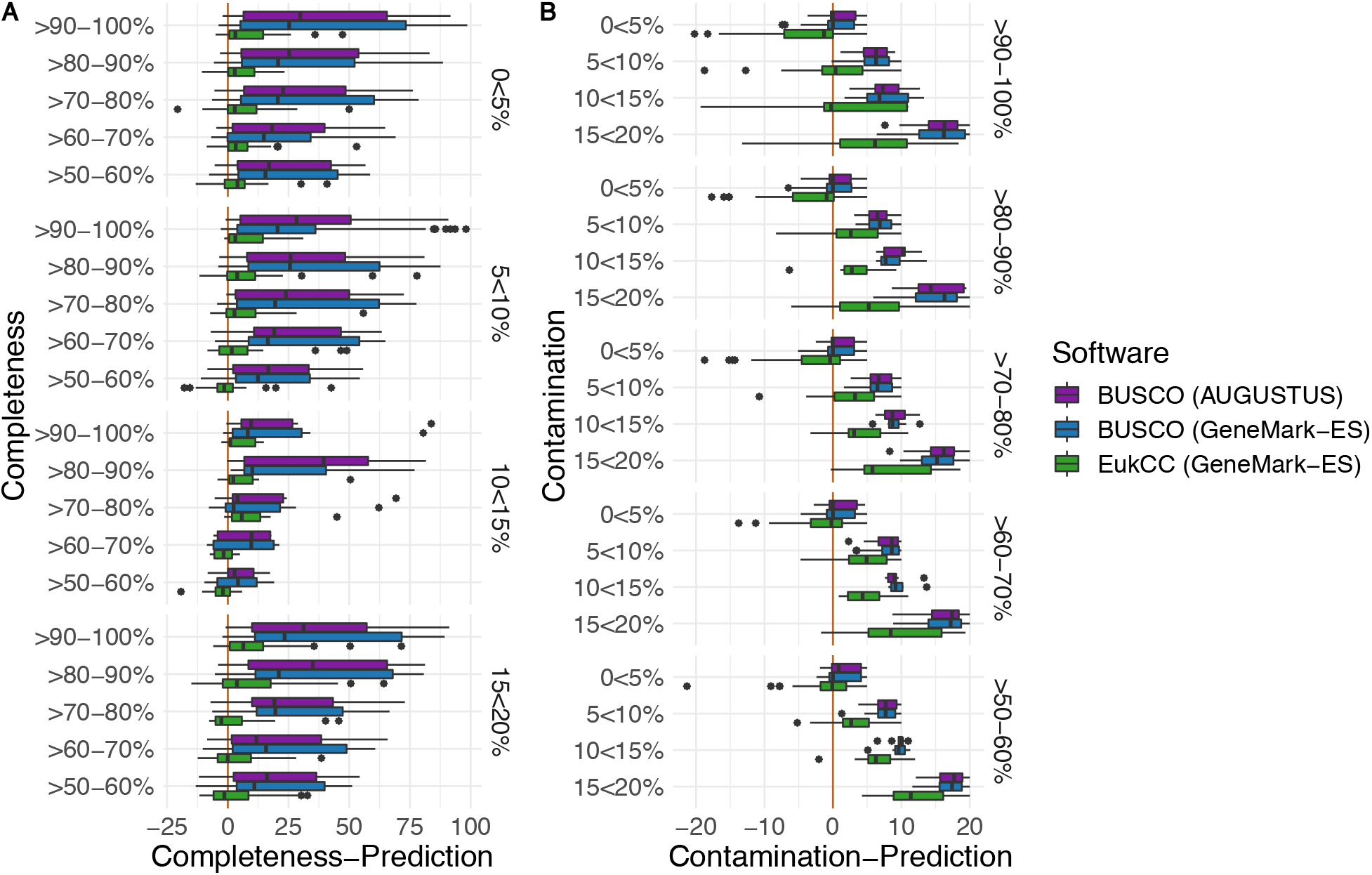
We compared EukCC to BUSCO using a set of 21 genomes from RefSeq belonging to alveolates, amoebozoa, apusozoa, fungi, rhizaria, stramenopiles and viridiplantae. We fragmented the genomes and added varying amounts of contamination from another genome in the same clade. We then ran BUSCO and EukCC to estimate completeness and contamination. The red line highlights zero percent deviation from the ground truth. **A)** We defined completeness in BUSCO as 100% minus missing BUSCOs. For genomes with a contamination between 0-5%, EukCC underestimated completeness with a median of 2.5%, while BUSCO underestimates the completeness across all genomes with a median above 20%. With increasing amounts of contamination, EukCC underestimates more rarely. Only when genome completeness falls below 50% and/or contamination exceeds 15% does EukCC consistently overestimate completeness. **B)** To evaluate contamination we counted the number of duplicated BUSCOs or marker genes (in the case of EukCC). For genomes with 0-5% contamination and high completeness (> 90%) EukCC overestimates contamination by below 5%. With increasing amounts of contamination, EukCC tends to underestimate contamination, but outperforms BUSCO, which consistently underestimates contamination by a larger fraction.

To demonstrate EukCCs performance in estimating contamination, we also assessed the contamination estimates against the known contamination rate at increasing levels of genome completeness (Figure 2 B, Figure S2). Contamination estimates were most accurate for fungi and Alvolates in genomes with completeness > 90% and simulated contamination < 5%, where EukCC deviates from the expected contamination estimate by less than 2%. Overall, EukCC tends to more frequently overestimate contamination compared to BUSCO. At lower levels of completeness (60-80%), EukCCs contamination estimates are less accurate, but as completeness increases (> 90%), the accuracy of contamination estimation increases, with a median error of < 2% for MAGs with contamination below 10%. Overall, as genomes include increasing amounts of contamination, EukCC begins to overestimate completeness, e.g. by ~ 5% for Fungal MAGs with expected completeness 60% < *x* < 70% and a contamination of 10% < *x* < 15% (Figure S2). This is somewhat to be expected, as there is a greater chance of finding an expected marker gene in the contaminating contigs, leading to inflated completeness. To investigate this, we added contamination in the form of random DNA to the MAGs and again estimated the completeness and contamination. In this case the contamination estimate is not affected by the added contamination, confirming the hypothesis as to the source of the overestimate.

To demonstrate that the EukCC SCMGs within this evaluation are distributed evenly across the entire genome, we randomly sampled 5 kb fragments and computed the Pearson correlation between the sampled size and the recovered marker genes for all species used within this benchmark. All sets used in this test showed linearity with a Pearson correlation coefficient of at least 0.95, indicating a uniform distribution of the marker genes across the genome.

As we could see a difference between BUSCO’s performance when using GeneMark-ES compared to AUGUSTUS, we investigated how well the GeneMark-ES predicted proteins overlap with annotations from RefSeq. For a taxonomically balanced subset of 89 eukaryotic genomes, we predicted proteins *de novo* using GeneMark-ES and cross referenced SCMGs used by EukCC against RefSeq annotated sequences from the same species using DIAMOND (Buchfink et al. 2015; (see Methods). We then generated a pairwise alignment between the predicted (query) protein and the best hit from the reference set and counted the gaps (irrespective of length) in both the reference and the query. Pairwise alignments with few gaps generally involve proteins of the same length. In the relatively few cases where there were a larger number of gaps (>10 gaps), these were introduced because the GeneMark-ES proteins were smaller compared to RefSeq, suggesting that GeneMark-ES does miss a small subset of exons. Despite this, the assigned RefSeq proteins and the corresponding GeneMark-ES proteins were found to have a generally similar length distribution. Together this suggests that the SCMGs chosen by EukCC and predicted by GeneMark-ES are similar to the annotations in RefSeq (Figure S4).

Across all simulated genomes, EukCC could estimate genome quality starting from a completeness of around 50 percent. Genomes less complete than this, were often not able to be processed using the self training mode of GeneMark-ES. In addition, GeneMark-ES failed to predict proteins for two Cryptophyta species, which were excluded from the benchmark. BUSCO with AUGUSTUS predicted an overall completeness of 3 and 3.6% for these genomes.

In this benchmark we found that BUSCO tends to underestimate contamination in genomes of high completeness (Figure S2) and underestimates completeness across all tested clades, except fungi. Meanwhile, EukCC tends to underestimate completeness and overestimate contamination (albeit at low rates), which leads to more conservative, yet more accurate genome quality estimates.

### Application of EukCC for the evaluation of MAG quality

Having established the utility of EukCC on the simulated benchmark, we applied it to metagenomic datasets. As a first example, we investigated samples from the skin microbiome, a relatively well characterised microbiome, where the community has low diversity and is known to include many Fungal species, many of which have been isolated and their genomes sequenced (Byrd et al. 2018; Wu et al. 2015). These features provided the best chance of producing *de novo* assembled eukaryotic MAGs for which we could estimate the quality using EukCC and independently verify their quality using reference genomes. Furthermore, given that BUSCO performs well for fungal genomes, this would provide additional validation of the EukCC results.

We retrieved the sequencing data for the largest publicly available human skin microbiome study (accession PRJNA46333, Oh et al. 2014; Oh et al. 2016), which comprises ~ 4,000 individual sequencing runs, from which 1483 runs can be assigned to 15 individuals. Following assembly with metaSPAdes (Nurk et al. 2017) and binning with CONCOCT (Alneberg et al. 2014; see Methods), 1573 of the assembled runs produced bins, generating 33,879 bins in total. As these bins were expected to be a mixture of bacterial and eukaryotic genomes (Findley et al. 2013; Tsai et al. 2016), a top level classification was performed of all bins using EukRep (West et al. 2018) to identify any bin containing at least 1 Mb of predicted eukaryotic DNA, reducing the number of bins from 33,879 to 279 (with the bins hereafter referred to as a MAG).

Using EukCC we could predict the MAG quality for 109 out of the 279 MAGs. We then assigned reference genomes to as many MAGs as possible, by finding the closest GenBank entry for each based on Mash distances (Ondov et al. 2016; (see Methods). 95.4% of the MAGs (104 out of 109) could be assigned to a fungal reference genome with a Mash distance < 0.1, corresponding to average nucleotide identity (ANI) of ~ 90% or above. We compared the alignment fraction of the reference to the predicted completeness of EukCC for all MAGs. For those MAGs that could be aligned to a reference genome with an ANI > 95% and had a predicted contamination below 5% and a completeness > 50%, the median difference between alignment fraction and predicted completeness was 3.6% (Figure S3 B).

We then computed completeness estimates for all MAGs using BUSCO (fungi set) and FGMP (another Fungal genome quality estimator) (Cissé and Stajich 2019). Using the genome based completeness estimation from the genome alignments, described earlier, as a true estimate of completeness for each MAG, we compared this to the corresponding completeness estimates from each of the three tools. BUSCO and EukCC assigned similar values of completeness to each MAG, while FGMP had a wider spread: FGMP overestimated completeness with a median of 16.2%, BUSCO underestimated completeness with a median of 8.5%, while EukCC showed the lowest deviation, underestimating completeness with a median of 3.1% (Figure S3 D).

Next, we removed the redundancy between the MAGs (based on the assignment to the same reference genomes and retaining the most complete MAG, with a contamination < 5%), This yielded a non-redundant set of 5 MAGs, corresponding to *Malassezia restricta* (with a EukCC reported completeness of 92.84% and a contamination of 1.38%), *M. globosa* (completeness 83.43% and contamination 2.25%), *M. sympodialis* (completeness 85.56% and contamination 0.79%), the unclassified *M. sp*. (completeness 83.05% and contamination 2.23%) and *M. sloof-fiae* (completeness 81.21% and contamination 2.37%). The average nucleotide identity (ANI) to the respective reference genome was above 98% for all MAGs but *M. sp*. (ANI 93.9%).

We found two additional MAGs that we could not assign to any known *Malassezia* species, but were identified by EukCC as likely to belong to the *Malasazzia* genus. We computed the mash distance between both MAGs and determined that they belong to the same unknown species (mash distance of 0.004). After dereplication, the representative MAG was estimated to have a completeness of 87.71% and a contamination of 1.18%. Wu et al. (2015) reported that Malassezia, in contrast to other Basidiomycetes, should contain the gene family that matches the Pfam entry DUF1214 (Pfam accession PF06742). We could verify the presence of this gene family in all reference Malassezia genomes except *M. japonica* and *M. obtusa*. We could also find this gene family in the MAGs assigned to *M. restricta, M. globosa, M. sloofia* and *M. sympodialis*, but not in the MAG lacking a species match nor in the MAG assigned to *M. sp*.. As both MAGs are predicted to be incomplete, this protein family could be missing by chance or due to misclassification of the MAGs. To verify if the lineage estimation from EukCC, assigning all MAGs to *Malassezia* genus, as well as the mash assignment, we identified SCMGs present in all Malassezia as well as in *Saccharomyces cerevisiae*, *Piloderma croceum* and *Ustilago maydis*. We used members of these protein families to build a tree that included all recovered non-redundant MAGs and all representative genomes from the Malassezia clade, as well as the aforementioned fungi. In the resulting tree, all MAGs cluster next to or close to their assigned reference genome. The tree recapitulates the three cluster structure first described by Wu et al. (2015). The MAG representing an unknown species is located within the Malassezia clade, and might be a member of clade B, confirming the taxonomic assignment by Mash and EukCC (Figure 3 A).

**Figure 3:**
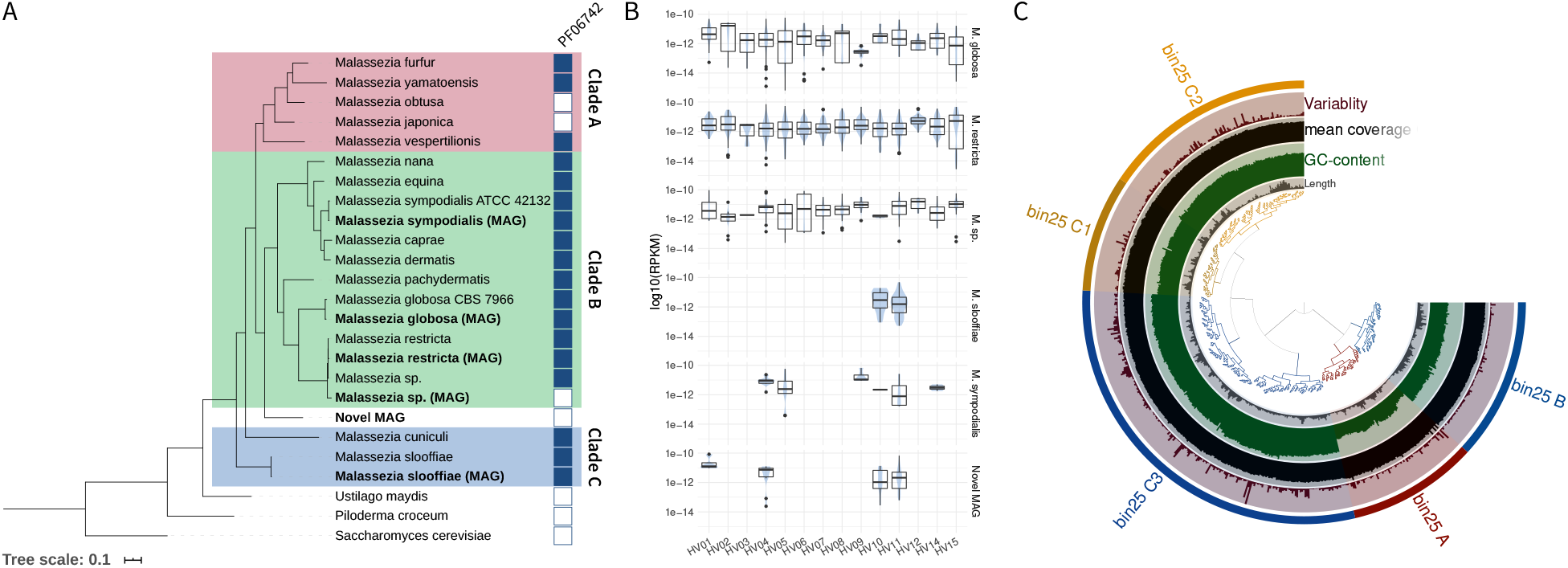
We assembled 1573 metagenomes and could recover almost complete MAGs of *M. globosa, M. restricta, M. sp., M. sloofiae* and *M. sympholidalis*. Additionally we recovered a *Malassezia* MAG with no known matching species. **A)** Using four genes occurring in single copy in all representative *Malassezia* species, in the recovered MAGs as well as in *S. cerevisiae* and two species of *Basidiomycota*, we constructed a phylogenetic tree with MAFFT and FastTree2. The tree recapitulates the clustering suggested by Wu et al. (2015), consisting of three clusters A, B and C. All recovered MAGs cluster next or close to their assigned species (bold). The MAG representing the unknown species (green) is clustered within the *Malassezia* clade, confirming the previous annotation. **B)** For each MAG we counted the RPKM if more than 30% of the genome was present in a sample. Using this approach, we could detect *M. globosa*, *M. restricta* and *M. sp*. across all individual subjects. The less prevalent *M. sloofiae* and *M. sympholidalis* could only be found in 2 and 6 individuals, respectively. The novel MAG could be found in four subjects. **C)** We analysed the MAG using anvi’o’s refine method. The clustering suggests a splitting into two main clusters A+B and C. While A has a lower GC content than the other clusters, all three clusters could be annotated as Malasezzia using UniRef90.

To investigate the prevalence of the five recovered MAGs, we aligned the reads from 1483 skin metagenomes belonging to 15 individuals to the MAGs and computed the Reads Per Kilobase of transcript per Million mapped reads (RPKM) of unique reads for samples if 30% of the target MAG was covered. Using this approach, we identified *M. globosa*, *M. sp*. and *M. restricta* in all individuals of this study (n = 15). The novel Malassezia species was present in 4 different individuals, which was more prevalent than *M. sloofia* (n = 2) and close to the prevalence of *M. sympodialis* (n = 6) (Figure 3 B).

We then inspected the potentially novel *Malassezia* species genome using anvi’o *refine* (Eren et al. 2015) and identified three contig clusters (Figure 3 C). Each subcluster was taxonomically analysed using matches to Uniref90 and could be associated to the genus *Malassezia* with a majority vote of at least 60% of the sampled proteins (see Methods). We also looked at the density of marker genes in each subcluster. With a density of 14.8% completeness per Mb DNA, subcluster C contributes 72.8% of absolute completeness and is the most marker gene rich cluster, compared to B (8%/Mb) and A (5%/Mb). While cluster A contigs have a lower GC content than the other anvi’o clusters and a lower density of marker genes, the taxonomic profile still suggests that it belongs to the genus *Malassezia*. Despite the differences in GC content and gene density, we decided to keep cluster A in the final MAG based on the consistency of the taxonomic assignments. However this cluster only contributes to 3.7% of the total completeness of the genome, so if it were to be omitted this would still represent a largely complete genome.

### Applying EukCC to a *Bathycoccus* MAG from TARA Ocean data

Having established that EukCC quality estimates were accurate in a well characterized community, we then tested it on samples in which we expect a diverse range of eukaryotes, beyond fungi. To do so, we focussed on the eukaryotic enriched samples (size fractionated samples in the range 0.5 μm to 2 mm (Protists size fraction, study: PRJEB4352)) from the TARA Oceans project (Carradec et al. 2018). As a prelude to investigating eukaryotes from this biome we randomly selected 10 out of the 912 available runs.

We assembled the samples using metaSPAdes and binned the resulting contigs using CONCOCT. After screening for eukaryotic bins using EukRep, we ran EukCC in default mode. Among the bins associated with ERR1726523, we identified a 13 Mb bin that EukCC estimated to have a completeness of 87.62% with a contamination of 0.32%. EukCC inferred a taxonomic placement in the order *Mamiellales* (green algae). We compared this MAG to eukaryotic genomes in GenBank using Mash, and found the closest match to *Bathycoccus sp. TOSAG39-1* (GCA_900128745.1, 10 Mb), with a Mash distance of 0.04. The taxonomy of this genome confirmed the EukCC inferred lineage and chosen SCMG set. We then aligned the MAG to this reference using dnadiff: in a pairwise alignment 52.01% of the MAG covered 78.97% of the reference genome with an ANI of 96.08%. The identified reference genome was published by Vannier et al. (2016) by merging four single-cell amplified genomes (SAGs). Vannier et al. (2016) estimated their SAG to be 64% complete using eukaryotic core genes from CEGMA. Using BUSCOs chlorophyta set we estimated the SAG to be 47.4% complete, with 4.8% marker genes duplicated. EukCC estimates the SAG to be 59.65% complete, slightly lower than the original estimation but higher than that suggested by BUSCO. However, EukCC indicated 14.04% contamination, which may have resulted from the merging of the SAGs.

The reported MAG has a scaffold N50 of 18.534 KB (contig N50: 10,754) and a scaffold size of 13.1 MB (1,097 scaffolds, 1,703 contigs). This compares favorably against TOSAG39-1 which has a N50 of 14,082 (contig N50: 13,604) and a scaffold size of 10.1Mb (2,118 scaffolds, 2,398 contigs). We evaluated the new MAG using BUSCOs chlorophyta set, which suggested the MAG to be 65.6% complete with only 4 BUSCOs duplicated (0.2%). While this estimate is 22% lower than EukCCs proposed completeness of 87.62%, it still shows an improvement of at least 10% and a notable reduction in contamination compared to the published *TOSAG39-1* genome. To check for assembly and binning errors, we again analysed the MAG using the bin refinement method in anvi’o (Figure S5): the anvi’o clustering divides the bin into two main clusters. Both clusters share similar GC content and coverage. From each cluster we inferred the taxonomic annotation by comparing a subsample of up to 200 proteins against Uniref90. For all analysed clusters the consensus lineage ended at the genus *Bathycoccaceae*, indicating a consistent MAG with no significant contamination.

## Discussion

Microbial eukaryotes represent a largely unexplored area of biodiversity. The use of modern genomic and metagenomic approaches are beginning to provide access to the genetic composition of these hitherto unknown organisms. However, in this study we have demonstrated that widely used tools for estimating eukaryotic genome quality (completeness and contamination) do not work uniformly across all microbial eukaryotes, which limits their application – for example within metagenomic pipelines.

Our results also highlight that the quality of the gene prediction step influences the quality estimates given by BUSCO – using NCBI Ref-Seq annotations instead of AUGUSTUS gene predictions raised the predicted average quality of the tested genomes. However, regardless of the gene annotations used, BUSCOs eukaryota set consistently underestimated genome completeness within certain clades. This within-clade error can not be explained by low quality reference genomes, but rather is indicative of a suboptimal eukaryota set. Thus, we showed that using a more specific marker gene set can lead to a better estimate, but BUSCOs protist set still did not lead to desirable results.

To overcome many of these limitations, we have developed EukCC, a novel tool to estimate microbial eukaryotic genome quality. EukCC uses a reference database to dynamically select the most appropriate out of 477 single copy marker genes sets. This set is then used to report genome completeness and contamination, as well as a taxonomic placement. Using simulated data, we showed that EukCC estimates genome quality across several taxonomic clades and performs on a par with, or better than, BUSCO. We showed that EukCC typically underestimates completeness and overestimates contamination. This conservative approach ensures that MAGs confirmed by EukCC are likely to be of high quality. EukCC also works independently of user input and can thus be used to analyse potential eukaryotic genomes from unknown species.

Nevertheless, we see a connection between the number of known species in a taxonomic group and the performance of EukCC. Some eukaryotic clades have very few high quality reference genomes. For example, at the time of writing, Apusozoa, Rhizaria, Cryptophyta as well as Rhodophyta, each have less than 10 reference genomes. While the current version of EukCC is known to perform better for more deeply sampled clades, we have demonstrated that the general framework can deliver consistent and high quality estimates across a broad taxonomic range. Thus, we aim to update the database regularly in order to build on growing public data and improve our performance across all clades.

Using EukCC, we are now able to systematically screen large libraries of previously ignored or unanalysed bins from published shotgun metagenomes. We showed that reanalysing published skin metagenomes we could find a novel species prevalent in ~ 25% of the analysed subjects. The novel species belongs to the well sampled *Malassezia* genus and could prove interesting in the context of understanding the skin microbiome. We have additionally demonstrated that current metagenomic techniques are also able to recover large fractions of eukaryotic genomes from more complex biomes, such as marine environment.

## Conclusion

With EukCC, we present an easy to use tool to estimate genome quality metrics for microbial eukaryotes and have demonstrated a substantial improvement in the applicability of EukCC compared to other tools. While this tool was developed with application to MAGs in mind, we do not see any limitation within EukCC to prevent it from being applied to SAGs, or even isolate genomes. To demonstrate the applicability of EukCC, we have identified two novel eukaryotic genomes from metagenomic samples, and have subsequently verified the quality of these genomes using a variety of approaches. EukCC provides the first step of many to assess the quality of MAGs and offers a way to select those that are likely to represent high quality MAGs.

## Methods

### Evaluation of BUSCO results

To evaluate BUSCO, we downloaded genomes and corresponding annotations for 418 eukaryotic reference species from RefSeq (Sep 26th 2019), excluding those belonging to bilateria or vascular plants. Each genome was annotated using GeneMark-ES (parameters: `‘-v -fungus -ES -cores 8 -min_contig 5000 -sequence input.fa’`). We then ran BUSCO (version 3.1) in using the ‘eukaryota_odb9’ BUSCO set in ‘genome mode’ using AUGUSTUS (version 3.3.2) as well as in ‘protein mode’ for both the RefSeq annotated proteomes as well as the GeneMark-ES predicted proteins. This procedure was then repeated for the ‘protist_esemble’ BUSCO set. To compare the BUSCO results to EukCC, we defined completeness as 100% minus the fraction of missing BUSCOs, and contamination as the fraction of duplicated BUSCOs. For Figure 1B, all reported BUSCOs in all analysed genomes were displayed using ComplexHeatmap (Gu et al. 2016) in R 3.5.1 (R Core Team 2018) and clustered the rows using ‘klaR’ (Weihs et al. 2005).

### GeneMark-ES *de novo* protein prediction comparison

Following this, we compared RefSeq provided annotation and proteins predicted from GeneMark-ES (parameters: ‘`-v -fungus -ES -cores 8 -min_contig 5000 -sequence input.fa’`). For each genome, we ran the BLASTp option from DIAMOND (Buchfink et al. 2015) on the proteins used by EukCC to estimate genome quality matching against the RefSeq annotated proteome of the same species. For the best hit, we aligned both sequences using MAFFT (Katoh et al. 2002). Subsequently, we compared the length distributions between GeneMark-ES and RefSeq annotated sequences. Additionally, we counted the number of gaps within the alignment occurring in either the reference or the query sequence. Analyses were performed using R 3.5.1 and plots were generated using ggplot (Wickham 2016).

### EukCC reference database creation

To build EukCC’s database we downloaded the genomes of 754 eukaryotic species from NCBI GenBank and RefSeq, all of which were either marked as representative genomes (August 1st 2019) or used as UniProt reference proteomes (May 28th 2019) (“UniProt” 2019) excluding those belonging to bilateria or vascular plants (Table S1). Following this, we predicted the proteome of each genome using GeneMark-ES and annotated the resulting proteins using PANTHER families 14.1 with hmmer 3.2.1 (Figure S6 A)). During this process 20 genomes were excluded due to GeneMark-ES failing to produce an output, reducing the number of species to 734. The failure was mostly caused by fragmented reference genomes, making it impossible for GeneMark-ES to pass the training step.

Subsequently, using the annotated proteins of a taxonomically balanced subset of species, we defined bit score gathering thresholds for each PANTHER profile HMM. For this, we chose at most 30 genomes per major sub-clade of eukaryota (e.g. Opisthokonta, Amoebozoa, Alveolata) and sampled evenly across all phyla below. Within these species, we identified the bit score value maximizing the number of single hits for each profile HMM.

After applying the bit score thresholds across all annotated species, we searched for profiles covering all species as single copy markers. As no single copy markers spanning all species could be found, we used a greedy algorithm to define a reference set of overlapping single copy marker genes. The resulting reference set contained 55 profiles, covering each species within the training set as a single copy marker between 3 and 34 times. The single copy proteins belonging to each profile HMM within the reference set were aligned using MAFFT and horizontally concatenated. Consequently we used this alignment to build a reference tree using FastTree2 with default settings (Price et al. 2010).

Following this, we identified 477 sets of single copy genes with a single copy prevalence cut-off of 98% in each clade of at least 3 species.

### Overview of the EukCC algorithm

As a first step, EukCC uses Genemark-ES to predict proteins in the input genome (Figure S6 B). The EukCC pipeline then performs a two stage analysis to determine the best set of SCMGs for downstream analysis. The first stage uses the reference set to define a first approximate taxonomic classification of the MAG to enable the placement in the precomputed reference tree using pplacer version v1.1.alpha19 (Matsen et al. 2010). For each protein, the best placement as indicated by the posterior likelihood is chosen. Using these placements, EukCC relies on ete3 to compute the lowest common ancestor (LCA) or the highest possible ancestor (HPA) for which a set of single marker genes exist (Huerta-Cepas et al. 2016). In a second stage, the HMMs defined in the chosen SCMG set are scanned against the predicted proteome using hmmer. The fraction of existing profiles is reported as completeness, and the fraction of duplicated markers is reported as contamination. Finally, EukCC reports a lowest common ancestor lineage of the input genome, based on the species within the marker set.

### Evaluation data creation

In order to benchmark EukCC and BUSCO with known data, we created in silico fragmented and contaminated genomes. For this we chose RefSeq genomes across all relevant taxonomic clades, which were not included in the initial training data. From each clade we selected up to 4 species to evaluate completeness and contamination. If a selection of species could be made, we first included species from a rank not included in the training set, prioritising novel phyla over novel order and so forth. We then created fragments by stepping along chromosomes with step size chosen from a Poisson distribution (*rpois*(*n, λ* = 100) *×* 1000) and a minimum step size of 2000. Fragments were rejected or included at random to create a genome of a target size fraction. Contaminating contigs were sampled from different species from the same clade and were fragmented in the same way and combined to make a test genome.

### Benchmark and comparison of EukCC to BUSCO

Following the creation of the benchmark data, we ran BUSCO (version 3.1) in ‘genome mode’ using the AUGUSTUS gene predictor (version 3.3.2) on the simulated genomes. For each genome we used the most suitable set of BUSCOs for the data. For example, when assessing a protist genome, we used the ‘protists_ensembl’ set, and for fungi we used the universal ‘fungi_odb9’ set. Notably, we used the protist set to evaluate the alveolata species, as BUSCOs performance decreased when using the more specific ‘alveolata_stramenophiles_ensembl’ set. We then used the ‘short_summary_*’ files from which we extracted the percentage of missing and duplicated marker genes. We defined completeness as 100% minus the percentage of missing BUSCOs, thus also including fragmented BUSCOs in the completeness score. Additionally, we ran BUSCO in protein mode using proteins predicted by GeneMark-ES (parameters: `‘-v -fungus -ES -cores 8 - min_contig 5000 -sequence input.fa’`). Eu-kCC was used with default parameters and database version 1. We discarded any simulated MAG from the benchmark for which not all three methods could produce a quality estimate. Finally we obtained 678 results per evaluated algorithm, which were aggregated with R using dplyr and plotted using ggplot.

### Assembly and binning of skin metagenomic datasets

We downloaded 3,963 shotgun metagenomic datasets from the skin metagenome study PRJNA46333. Each dataset as assembled using metaSPAdes (version 3.12, default parameters in metagenomics mode [`-meta`]) (Nurk et al. 2017) and binned using CONCOCT (version 1.0) (Alneberg et al. 2014) as part of the metaWRAP (version 1.1) (Uritskiy et al. 2018). Subsequently, we estimated the genomic composition in each bin using EukRep (version 0.6.5) and bins with more than 1 Mb eukaryotic DNA were selected for further analysis. Selected ‘eukaryotic’ bins were then analysed using EukCC and compared to RefSeq and GenBank (both retrieved Sep. 26 2019) entries by comparing Mash distances (version 2.2.2 default parameters) (Ondov et al. 2019) and subsequently using dnadiff (from the mummer package, version 3.23) (Kurtz et al. 2004) for the top hit, if the Mash distance was below 0.1.

### Tree building and analysis of skin MAGs

To construct a phylogenetic tree for the 6 selected skin MAGs, 19 reference genomes of 16 Malasezzia species and *Saccharomyces cerevisiae*, *Ustilago maydis* and the GenBank entry of *Piloderma croceum*, we identified 4 SCMGs genes used by EukCC found in all genomes: PTHR10383, PTHR11377, PTHR12555, PTHR15680. Using MAFFT, in einsi mode, we aligned the protein sequences for each PANTHER entry before building a concatenated alignment file, which was used as input to FastTree2 to build the tree, using default settings. We visualized and rooted the tree using *S. cerevisiae* as an outgroup with iTOL v5 (Letunic and Bork 2016). Using hmmer 3.2.1 (`hmmscan -cut_ga`) we searched for the Pfam (El-Gebali et al. 2018) entry DUF1214 (Pfam accession PF06742, Pfam version 32) in the 6 MAGs and the 19 reference genomes. To further verify the quality of the MAG, we clustered the contigs using anvi’o’s refine module and sampled up to 200 proteins from each cluster. Each protein was compared against the UniRef90 database using DIAMOND (`parameters: blastp -threads 10`). Using the majority voted consensus lineage of up to three hits per protein (e-value threshold of 1e-20) with a majority threshold of 60% and a subsequent global majority vote using the same threshold, we assigned taxonomic lineages to each cluster.

### Analysis of TARA Ocean data

We assembled and analysed metagenomes from the TARA Ocean study PRJEB4352 using the same protocol as for the skin metagenomic data. We assembled and binned reads from ERR1700893, ERR1726523, ERR1726543, ERR1726560, ERR1726561, ERR1726573, ERR1726589, ERR1726593, ERR1726609, ERR1726612. The study we selected has 912 runs associated, and we chose this subset of runs at random as we were limited by the large amount of memory and CPU time required for each assembly (for example, assembling ERR1726589 required 942 Gb of RAM).

### Availability of data and materials

The EukCC code is available through Github (https://github.com/Finn-Lab/EukCC). Documentation can be found at readthedocs (https://eukcc.readthedocs.io/en/latest). The EukCC database can be downloaded from http://ftp.ebi.ac.uk/pub/databases/metageno-mics/eukcc_db_v1.tar.gz. All MAGs have been submitted into ENA under the accession PRJEB35744.

## Acknowledgements

Paul Saary is a member of Queens’ College, University of Cambridge.

## Supplementary material

**Figure S1:**
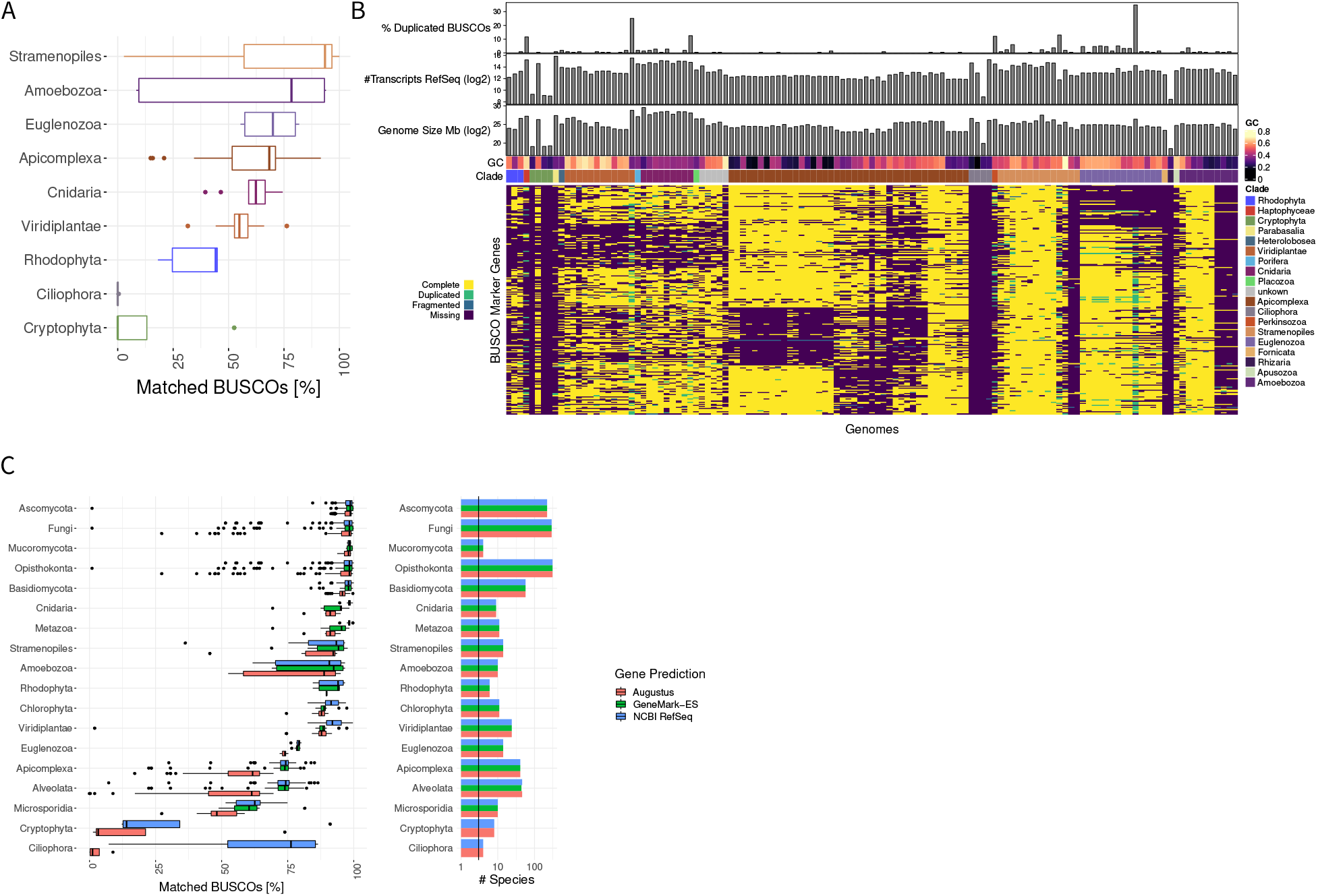
**A)**Corresponding to the analysis in Figure 1 we used the BUSCO protist set to analyse a set of non Fungal genomes. In a number of protist clades, e.g. Apicomplexa and Amoebozoa, not all BUSCOs can be found. **B)** Matrix showing the breakdown of missing BUSCOs across different taxonomic groups. **C)** Running BUSCO using the ‘eukaryota_odb9’ set in *genome mode*, using GeneMark-ES, or using the NCBI RefSeq annotations, the number of found BUSCOs across these three gene callers is similar for Fungal clades, but increases when using GeneMark-ES or RefSeq for Euglenzoa and Apicomplexa.

**Figure S2:**
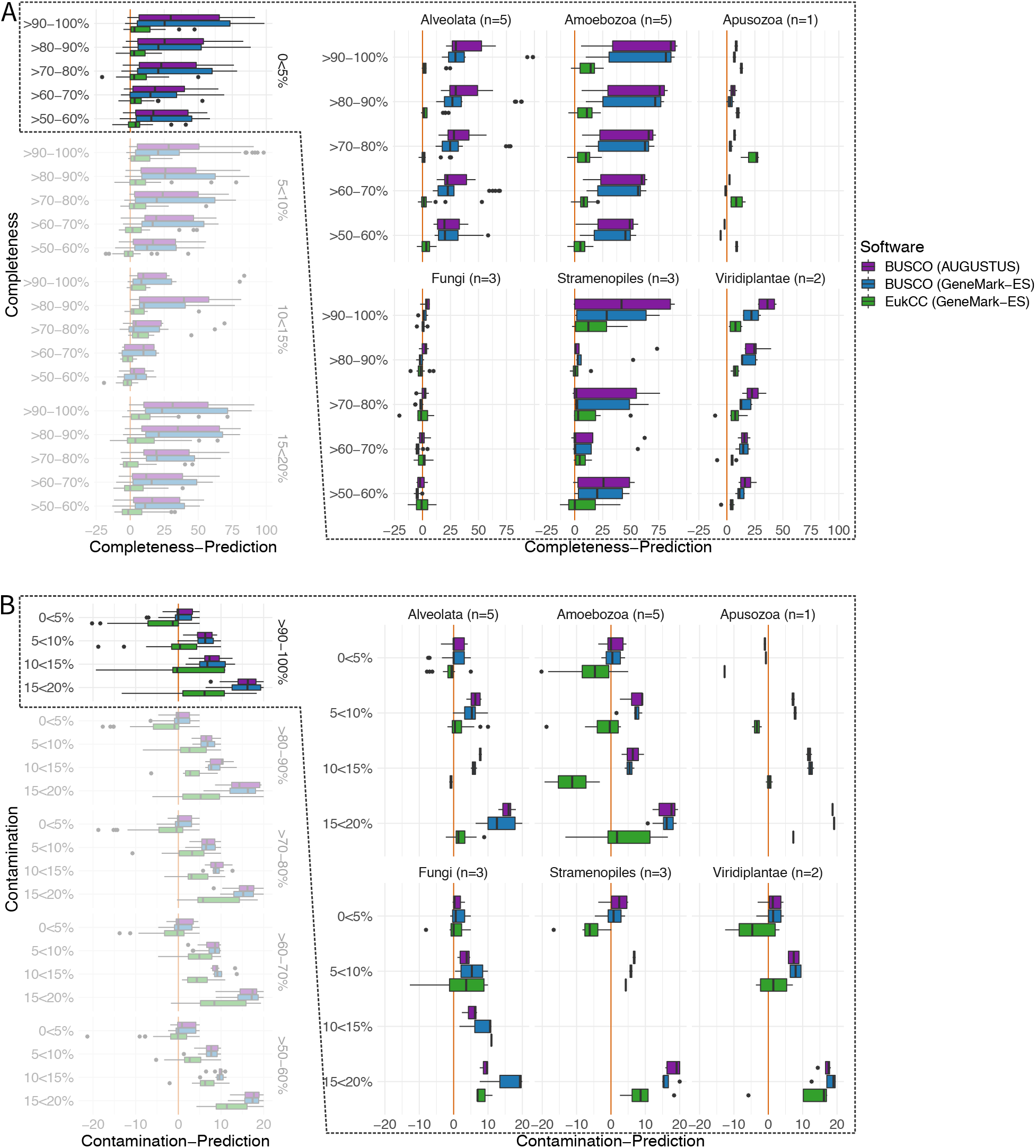
**A)** For simulated genomes with a contamination below 5% we split the panel 1 into the taxonomic clades (number of species per clade indicated by n). For the fungal clade both EukCC and BUSCO, independent on the gene caller, perform close to the expected value across a large range of simulated completeness. For Alveolates, Amoebozoa and Viridiplantae EukCC consistently performs better than BUSCO. Within the Stramenopiles EukCC both methods show a large variability in their performance. **B)** When looking at contamination estimated for highly complete genomes, BUSCO and EukCC perform best for low contamination ratios (<5%). For Alveolata EukCC performs well across a large range of contamination. In the fungal clade both BUSCO and EukCC perform better for low contamination ratio and start to underestimate contamination with increasing amount of contamination. Within Amoebozoa and Viridiplantae EukCC, in contrast to BUSCO, tends to overestimate contamination by aprox 5% for genomes with 0-5% contamination.

**Figure S3:**
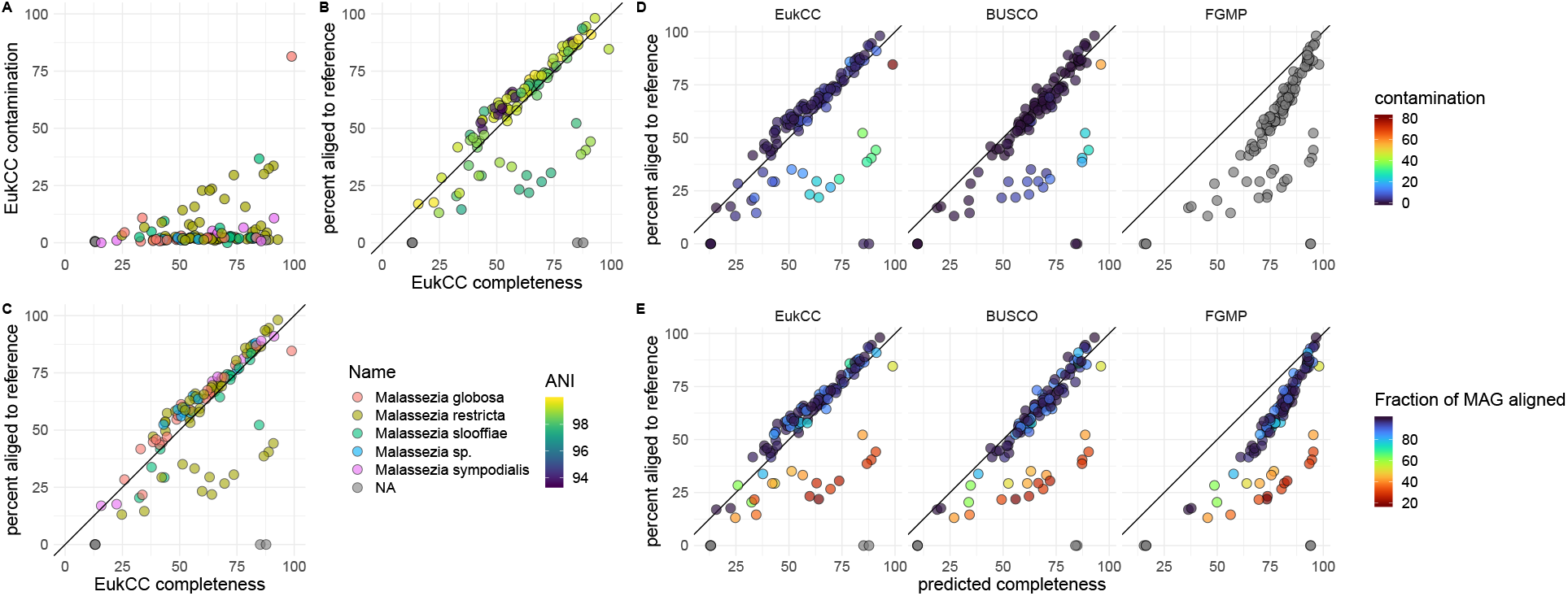
**A)** Bins recovered from skin metagenomes were assigned to a reference genome and then estimated using EukCC in terms of completeness and contamination. Bins could be assigned to five different species of *Malassezia*. **B+C)** For bins that could be assigned to a reference we compared predicted completeness to how much of the reference could be aligned to the bin. For most bins EukCCs prediction is close to the aligned fragment. We see no signal when comparing the prediction of EukCC to the average nucleotide identity (ANI) between the MAG and the assigned reference genome neither when color coded by assigned species. **D)** We thus also checked completeness using BUSCO and FGMP: BUSCO and EukCC performed comparable, both slightly underestimating completeness. FGMP overestimated completeness in almost all bins. Bins clearly overestimated by EukCC or BUSCO were also the most contaminated bins, which explains this behavior. **E)** When color coding bins by their percentage which could be aligned to the reference (Fraction of MAG aligned), well aligned bins are close to the diagonal and bins with a lower fraction of aligned DNA are commonly below the diagonal, which is in good agreement with the contamination estimate.

**Figure S4:**
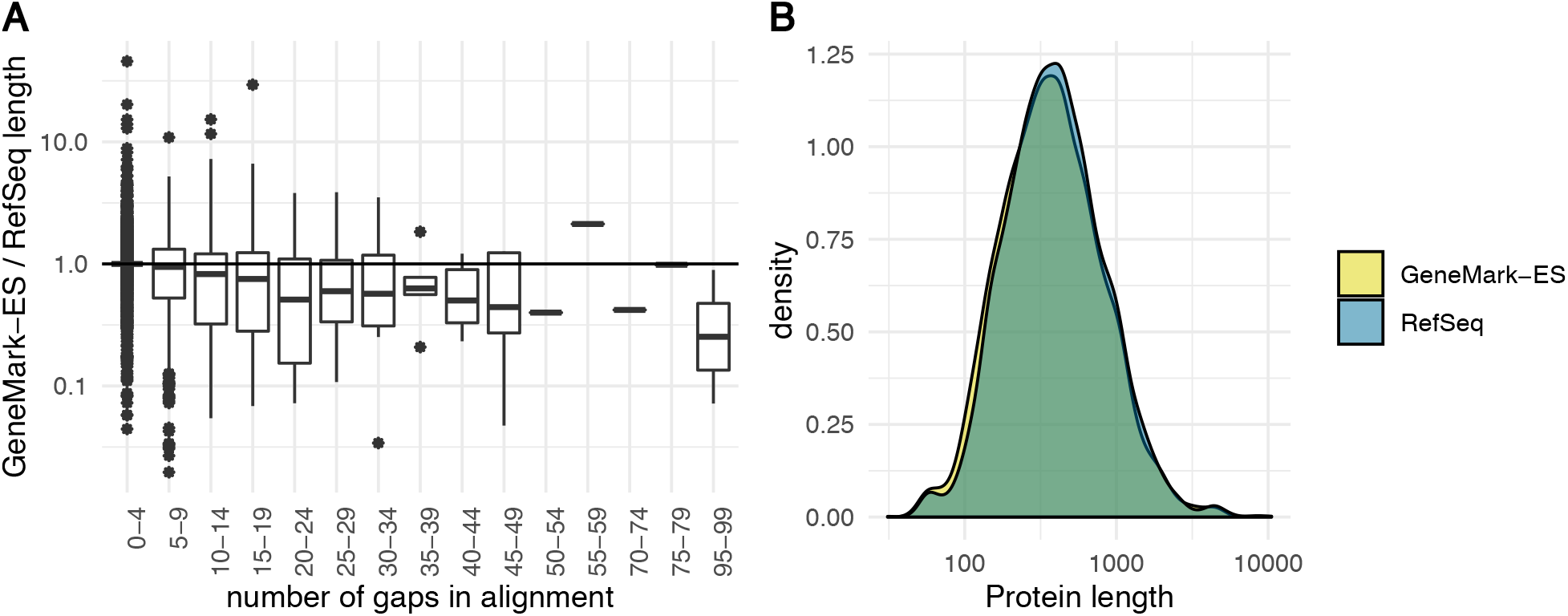
Proteins predicted with GeneMark-ES that occured in a single copy in representative genomes across the eukaryotic genome (see Table S2 for used genomes) were blasted against their RefSeq proteome. For each protein the best hit (judged by e-value) with a maximal e-value cutoff of 1e-5 was chosen as the corresponding true protein. **A)** Each protein was aligned to its reference protein using MAFFT and gaps were counted. With increasing number of gaps the predicted proteins are shorter than their reference, suggesting that GeneMark-ES seems to miss some introns. **B)** We could not find a systematic bias between the protein length of UniProt or GeneMark-ES predicted proteins.

**Figure S5:**
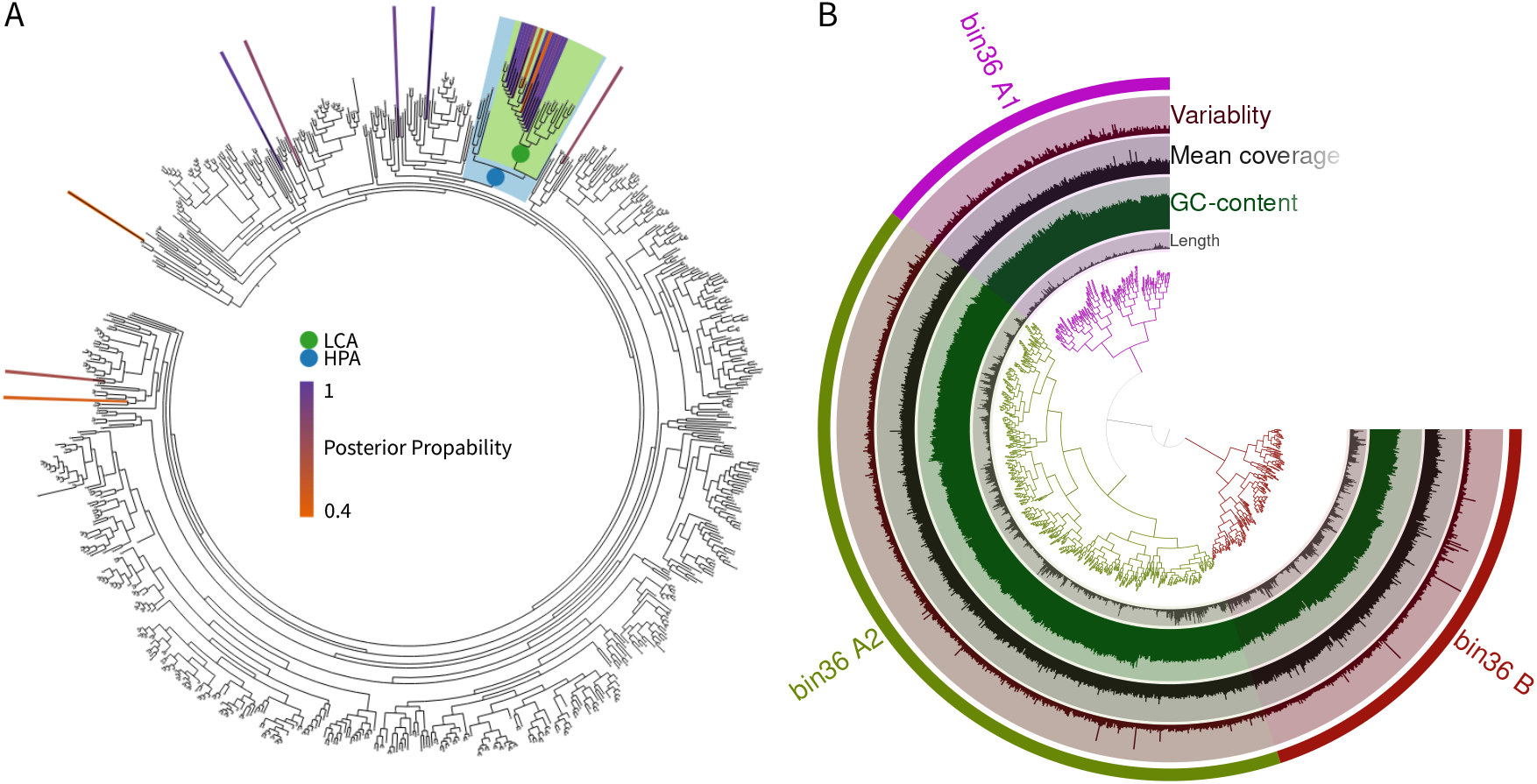
**A)** Using EukCC to estimate the quality of the reported *Bathycoccus* MAG, pplacer placed the found marker proteins from the reference set mostly within a small range of the phylogenetic tree. Some outliers were ignored by EukCC while choosing the LCA set (green). **B)** The recovered *Bathycoccus* MAG from the TARA Ocean data was checked for quality issues using anvi’o. While the GC content and the coverage across all contigs is very uniform, anvi’o could form two large clusters. We taxonomically analysed both clusters and the indicated subclusters. All groups could be assigned the same taxonomic lineage, suggesting low amounts of contamination.

**Figure S6:**
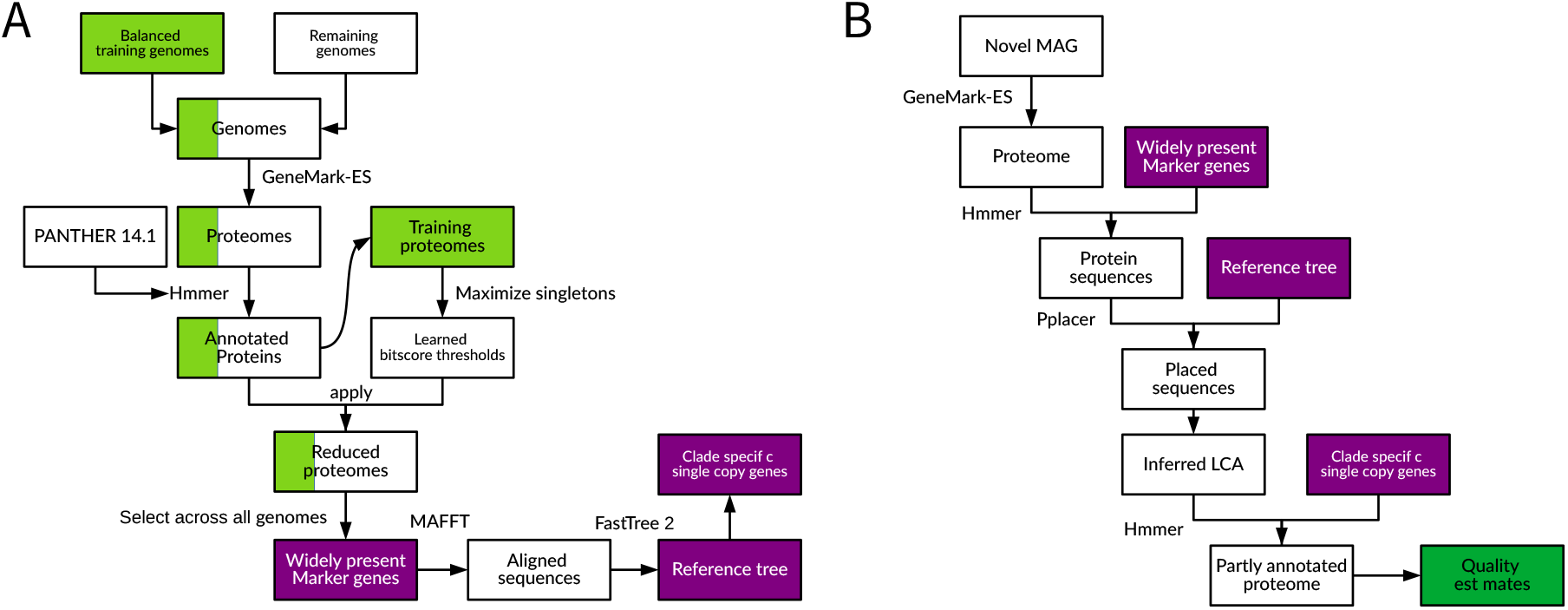
**A)** EukCCs database is created by predicting proteomes using GeneMark-ES first. Proteomes are annotated using hmmer with PANTHER 14.1 families. A predefined subset of proteomes (green) are then used to learn bitscore thresholds for all profile hmms, maximizing the singleton prevalence across this set. Annotations are then filtered using the thresholds. By choosing widely present single copy genes to cover the entire genome space several times, we build a tree by first aligning proteins independently and then concatenating the alignments. **B)** EukCC searches for the widely defined marker genes to used pplacer to place a novel MAGs proteins into the reference tree. Choosing the lowest common ancestor set quality is computed.

---

**Table S1:** Genomes used to train the database. Genomes were excluded because of GeneMark-ES failure (labeled: gmes), because of long branches in the tree (tree) or because of problems during the set creation (set).

**Table S2:** Genomes used to evaluate EukCC. Each simulated MAG was based on a single RefSeq entry with added fragments from the contaminant. The specified BUSCO set was used to evaluate and the clade used to group results for Figure S2.

**Table S3:** Looking at the novel MAG in anvi’o, we saw several clusters (Figure 3). For each cluster we calculated the size and the contribution in completeness as well as the average marker density. Cluster A has the lowest contribution as well as the lowest marker density. Cluster C1, C2 and C2 are similar in density and comprise the largest percentage of the MAG. Cluster B is between A and C in all measures.

